# Highly efficient homology-directed repair using Cas9 protein in *Ceratitis capitata*

**DOI:** 10.1101/323113

**Authors:** Roswitha A. Aumann, Marc F. Schetelig, Irina Häecker

## Abstract

**Background:** The Mediterranean fruit fly *Ceratitis capitata* is a highly polyphagous and invasive insect pest, causing vast economical damage in horticultural systems. A currently used control strategy is the sterile insect technique (SIT) that reduces pest populations through infertile matings with mass-released, sterilized insects. Transgenic approaches hold great promise to improve key aspects of a successful SIT program. However, there is strict or even prohibitive legislation regarding the release of genetically modified organisms (GMO), while novel CRISPR-Cas technologies might allow to develop genetically enhanced strains for SIT programs classified as non-transgenic.

**Results:** Here we describe highly efficient homology-directed repair genome editing in *C. capitata* by injecting pre-assembled CRISPR-Cas9 ribonucleoprotein complexes using different guide RNAs and a short single-stranded oligodeoxynucleotide donor to convert an enhanced green fluorescent protein in *C. capitata* into a blue fluorescent protein. Six out of seven fertile and individually backcrossed G_0_ individuals generated 57-90% knock-in rate within their total offspring and 70-96% knock-in rate within their phenotypically mutant offspring.

**Conclusion:** Considering the possibility that CRISPR-induced alterations in organisms could be classified as a non-GMO in the US and Europe, our approach to homology-directed repair genome editing can be used to genetically improve strains for pest control systems like SIT without the need to struggle with GMO directives. Furthermore, it can be used to recreate and use mutations, found in classical mutagenesis screens, for pest control systems.

## Background

With a large host range of more than 250 fruits, vegetables and nuts, a broad acceptance of both natural and cultivated habitats and tolerance over a comparatively wide temperature range the Mediterranean fruit fly, *Ceratitis capitata* (Wiedemann; Diptera: Tephritidae), has become one of the most successful invaders and thereby one of the most devastating and economically important insect pests worldwide [1–4].

In an attempt to reduce the use of insecticides in the fight against this and other crop pests, an effective, environmentally friendly, species-specific and area-wide control method has been the sterile insect technique (SIT) [5]. SIT is based on the mass release of sterilized male insects into a wild-type (WT) population, leading to infertile matings and thereby to a decrease in the number of progeny. Repeated releases thus allow for the suppression of a pest population to an economically uncritical size or to prevent the infestation of new areas.

There are several steps to be developed and improved to enable successful SIT programs. One is the generation of male-only populations also called *sexing*. Male-only releases are more effective and avoid the release of females that could still damage the fruits and crops by oviposition even if the eggs will not develop due to the sterilization step [13]. However, to date the release of transgenic organisms is highly regulated or even prohibited. Therefore, a tool that is able to create efficient and safe sexing systems, similar to classical mutagenesis and acceptable for a release, is needed.

In Europe, the deliberate release of genetically modified organisms (GMOs) is regulated by the ‘GMO Directive’ of the European Parliament and the Council on the deliberate release of genetically modified organisms into the environment (Directive 2001/18/EC, Council directive 90/220/EEC). Organisms covered by that directive have to undergo an environmental risk assessment to obtain authorization, and are subject to traceability, labelling, and monitoring regulations [14]. However, genetically modified organisms created via mutagenesis techniques are not included in the GMO Directive (‘the mutagenesis exemption’ of the GMO Directive of 2001, according to the EU court of justice). By its definition, mutagenesis involves the alteration of the genome of a living species but does not, unlike transgenesis, entail the insertion of foreign DNA into the organism [15].

Traditional mutagenesis techniques include chemical or UV mutagenesis, which both create random mutation products [19]. Either by the non-homologous end-joining (NHEJ) or the homologous-directed repair (HDR) pathway. While the NHEJ pathway is, in simplified terms, a rather ‘error-prone’ pathway, causing random insertions or deletions of nucleotides at the target site, the HDR pathway can be exploited to precisely manipulate the target sequence by providing a suitable DNA repair template including the desired alteration [20]. This allows the introduction of specific sequence changes without leaving exogenous DNA sequences in the genome. Therefore, once established in a new pest species, CRISPR-Cas HDR could be the long-awaited tool to overcome the disadvantages of conventional mutagenesis and transgenic methods to establish strains for SIT programs. This could improve the generation of insect strains without unintended and unknown changes in the genome caused by traditional mutagenesis.

To achieve specific genome alterations via CRISPR-Cas HDR in a highly effective manner, it is necessary to shift the equilibrium of NHEJ and HDR towards the less efficient HDR pathway [21]. At the same time, the balance between the two repair pathways differs widely among species and between different cell types of a single species as well as different cell cycle phases of a single cell [21]. Improving the efficiency of HDR was explored by the inhibition of key enzymes of the NHEJ pathway like the DNA ligase IV using the inhibitor Scr7 [22, 23] or the controlled timing of Cas9 delivery according to cell-cycle dynamics [24, 25]. Other important aspects for a precise HDR event are the prevention of re-editing of already modified loci, for example by introducing mutations in the protospacer adjacent motif (PAM) sequence or the guide RNA (gRNA) target site of the repair template, as well as considering the effects of the distance between the DSB site and desired mutation position on the mutagenesis efficiency [26, 27].

To determine the efficiency of such HDR-improving methods, a strategy for the simultaneous quantification of HDR and NHEJ events is targeting an enhanced green fluorescent marker protein (eGFP) and converting it into the blue fluorescent protein (BFP) [28]. This can be done by two single base substitutions in the chromophore of eGFP [28, 29]. Thereby, green fluorescence shows the absence of a mutation event, blue fluorescence indicates an HDR event and the loss of fluorescence represents unspecific mutation events caused by NHEJ repair. So far, in Medfly, mutant phenotypes could only be generated by NHEJ repair after CRISPR-Cas9-based gene disruption [30].

Here, we report the first and highly efficient CRISPR-Cas HDR knock-in of a short single stranded oligodeoxynucleotide (ssODN) repair template in *C. capitata*, by injecting *in vitro* preassembled Cas9-gRNA ribonucleoprotein complexes and a single-stranded oligo donor into *C. capitata* embryos carrying an eGFP marker gene to convert eGFP to BFP.

## Results

### Selection of gRNAs for eGFP mutagenesis in *C. capitata* and off-target analysis

Two previously evaluated guide RNAs against eGFP [28] were used to lead the Cas9 protein towards the chromophore of the eGFP marker gene of the transgenic *C. capitata* strain *TREhs43-hid*^*Ala5*^_F1m2 [31]. The gRNAs target the same region, therefore one HDR repair template (single-stranded oligodeoxynucleotide blue fluorescent protein, ssODN_BFP [28]) was used for both. They differ, however, in their orientation and cleavage site, as well as in the number of mismatches to their target sequence resulting from successful HDR (Fig. 1A) and their off-target activity.

**Fig. 1.**
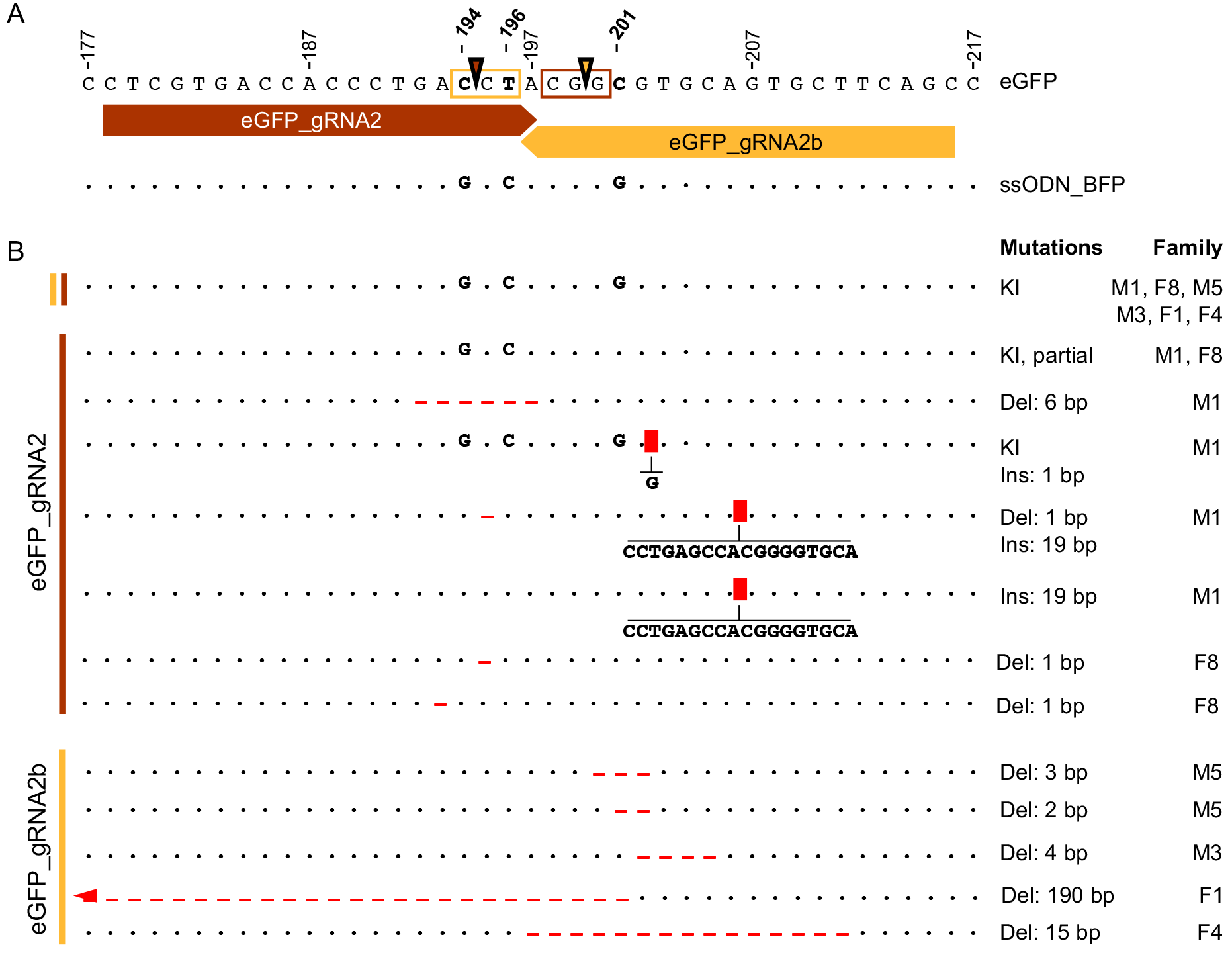
Position of gRNAs, protospacer adjacent motifs (PAM), double strand brakes (DSB) and single nucleotide polymorphisms (SNPs) within the eGFP target sequence. A) Relative to the eGFP sequence the eGFP_gRNA2 (red) is sense- and the eGFP_gRNA2b (yellow) is anti-sense-oriented. PAM sequences are highlighted within the eGFP sequence, DSB sites indicated by triangles. Related gRNA, PAM, and DSB site match in color. The ssODN_BFP sequence differs from the eGFP sequence in three positions, SNPs are (194C>**G**, 196T>**C**, 201C>**G**), and consensus is shown as dots. B) Sequences of mutant eGFP alleles identified in G_1_ individuals compared to the eGFP reference sequence. Explanation of indications and abbreviations: consensus is shown as dots, knock-in (KI) mutant sites in uppercase letters, deletions (Del) by red lines, insertion sites (Ins) as red rectangles. Families that were carried the respective mutation(s) are indicated.

*In silico* target site analysis predicted an on-target activity score of 0.272 for the eGFP_gRNA2 (scores are between 0 and 1; the higher the score the higher the expected activity [32]) and zero off-targets sites in the medfly genome (100% off-target score). eGFP_gRNA2b has an on-target activity score of 0.329 but two off-targets (98.94% off-target score: #1 score 4.23%; location NW_004524467.1 4,259 338 < 4,259,360; #2 score 1.13%; location NW_004523691.1 10,017,309 < 10,017,331; Ccap 1.1). Both off-target sites of eGFP_gRNA2b show four mismatches to the reference genome sequence. Importantly, none of the off-target sites are located in a coding sequence of *C. capitata* genome.

The repair template, ssODN_BFP, differs from the eGFP sequence by three bases (194C>G, 196T>C, 201C>G; Fig. 1A), whereby the first change (194C>G; Thr65>Ser65) causes a reversion of eGFP back to wild-type GFP and the second (196T>C; Tyr66>His66) converts GFP to BFP [28]. The third SNP (201C>G) is a silent mutation to further reduce the gRNA-target sequence similarity after HDR and thus prevent re-editing of the target sequence [28] (Fig. 1 A).

### CRISPR-Cas9 HDR mutagenesis in Medfly

Three injections were conducted to perform eGFP mutagenesis in the Medfly target line, homozygous for the eGFP marker gene. Two injections were performed with the gRNA eGFP_gRNA2 and one with gRNA eGFP_gRNA2b (Fig. 1 A). One of the eGFP_gRNA2 injections additionally contained the ligase IV inhibitor Scr7 in the injection mix. Each G_0_ adult survivor was screened for eGFP fluorescence to confirm the presence of the CRISPR target site, and individually backcrossed. Their offspring (G_1_) were screened for eGFP and BFP fluorescence.

First, the **eGFP_gRNA2** was injected with recombinant Cas9 protein and the ssODN_BFP donor template into 243 embryos of the strain *TREhs43hid*^*Ala5*^_F1m2, homozygous for eGFP. 16 reached the larval stage of which eight survived to adulthood. These (three males and five females) were individually backcrossed to *EgII* wild type virgin females and males, respectively. Eggs of these crosses were collected three times, at an interval of one to two days. Three crosses (M1, F2, F8) were fertile and two out of these three families produced phenotypically WT offspring missing the eGFP marker (Fig. 2). This effect was observed in 98 out of 116 flies (84%) in family M1 (Fig. 3 A), and 34 out of 42 flies (81%) in family F8 (Fig. 3 D). The loss of the eGFP fluorescence was interpreted as a positive CRISPR event (insertion/deletion or knock-in event at the target site). Blue fluorescence was not observed in any of the G_1_ flies.

**Fig. 2.**
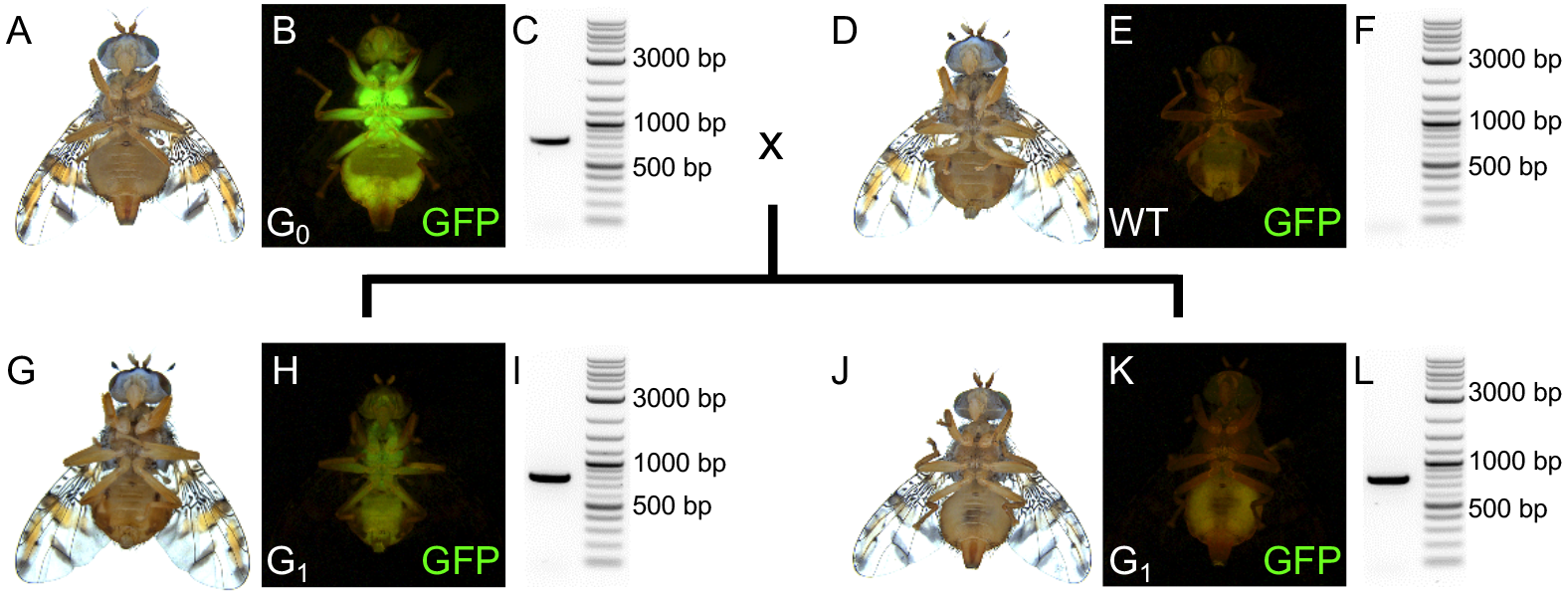
Crossing scheme and analysis of G_0_ and G_1_ individuals. Shown are fly images in bright field (A, D, G, J) and corresponding eGFP fluorescence (B, E, H, K) as well as the respective PCR validating the presence or absence of the eGFP marker gene (C, F, I, L). The *TREhs43hid*^*Ala5*^_F1m2 G_0_ individual, homozygous for the eGFP marker gene, injected with Cas9 and eGFP _gRNA2 or −2b, was crossed to WT *EgII* flies. G_1_ offspring was either heterozygous for the eGFP marker (H) and positive in eGFP-specific PCR (I), or phenotypically missing the eGFP fluorescence (K), but still carrying the eGFP marker gene (L), which indicates a CRISPR-induced mutagenesis. DNA ladder used for agarose gel is the 2log DNA-ladder (NEB); bp = base pair.

In a second, independent experiment, 323 embryos of the target line were injected with **eGFP_gRNA2 and 1 mM Scr7** additionally added to the injection mix. 79 reached the larval stage with 31 surviving to adulthood (17 males, 14 females). These were then backcrossed individually and eggs collected from 27 fertile crosses as described previously. In total, 1967 G_1_ offspring were screened for the loss of eGFP fluorescence. However, none of the families produced offspring phenotypically missing the eGFP marker.

Thirdly, **eGFP_gRNA2b**-Cas9 complexes together with ssODN_BFP donor template were injected into 371 embryos, yielding 19 larvae and 9 adult flies (five males, four females). Four of the nine individual crosses (M3, M5, F1, F4) were fertile and produced offspring mainly phenotypically missing the eGFP marker (79% to 100%), indicating a CRISPR-induced mutation (Fig. 3 G, J, M, P).

**Fig. 3.**
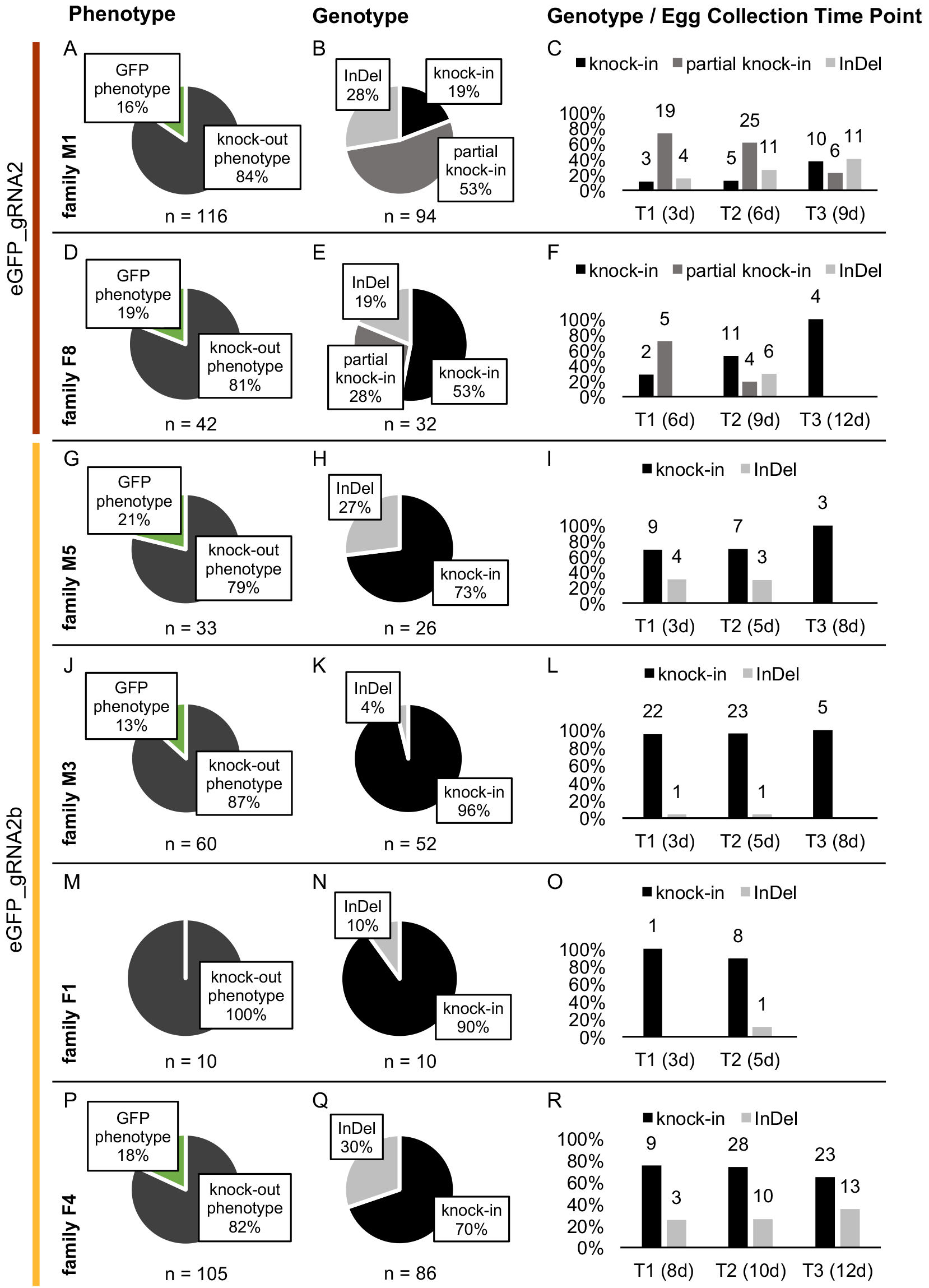
Frequency of CRISPR-Cas-induced G_1_ phenotypes and genotypes. Families M1 and F8 were injected with eGFP_gRNA2 (**A-F**), families M5, M3, F1 and F4 with eGFP_gRNA2b (**G-R**). In the first column, the absolute number of offspring per family and the occurrence of phenotypes “eGFP” (heterozygous) and “knock-out” (eGFP phenotypically missing) are shown (**A, D, G, J, M, P**). In addition, the second column shows the number of sequenced individuals with the frequency of different mutation types (knock-in, partial knock-in or insertion/deletion (InDel); **B, E, H, K, N, Q**). The third column shows the mutation types contingent upon egg collection time points (T1, T2, T3, (days after eclosion)) (**C, F, I, L, O, R**). Numbers above bars indicate absolute number of individuals per mutation per time point.

### Molecular verification of HDR or other mutagenic events

The genotype of all phenotypically WT G_1_ flies was analyzed via eGFP-specific PCR and subsequent sequencing, except for four individuals in family M1 and two in family F8, as DNA and sequence information could not be obtained for these flies.

Sequencing of DNA amplicons from individuals of **eGFP_gRNA2** injection revealed that 18 out of 94 phenotypically WT M1 offspring (19%) carried the complete knock-in genotype (three base pairs exchanged) and 50 (53%) carried a shorter version of the knock-in with only two of the three base pairs altered (194C>G, 196T>C). Both should lead to a loss of eGFP expression. The knock-out phenotype of the remaining 26 flies (28%) was caused by insertions or deletions in the target region (four different mutation events; Fig. 1 B, Fig. 3 B). In case of family F8, sequencing showed that 17 out of 32 phenotypically WT flies carried the complete knock-in genotype (three bp HDR) (53%) and nine carried the shorter version of the knock-in with two out of three bp mutated (28%). Moreover, two different deletion events caused by NHEJ repair were observed in six flies (19%: Fig. 1 B, Fig. 3 E).

Interestingly, the two different HDR events in G_1_ (complete three bp HDR versus two bp HDR) were not evenly distributed over the three egg collection time points (T1, T2, T3). In both families, the percentage of the complete HDR increased over time, whereas the rate of the partial HDR decreased. In the M1 family, 73% of the offspring from the first egg collection (T1) carried the partial knock-in, whereas only 22% of the offspring from the last egg collection (T3) carried this genotype. The complete knock-in was observed in 11.5% of the T1-offspring and in 37% of the T3-offspring of M1. In the F8 family, the partial knock-in decreased from 71.4% in T1 to none in T3. In contrast, the complete knock-in increased from 28.5% (T1) to 100% (T3) (Fig. 3 C, F).

In the second injection using **eGFP_gRNA2 plus Scr7**, no phenotypically wild type individuals were found during the screening and consequently no PCR amplicons could be generated or analyzed.

Analyzing the amplicons of the third injection with **eGFP_gRNA2b** confirmed efficient HDR in all four families, with 70% to 96% complete knock-in genotype within the phenotypically WT offspring. NHEJ caused one to two different mutation events per family, explaining the knock-out phenotype of the remaining flies (Fig. 3 H, K, N, Q; Fig. 1B). The occurrence of HDR events increased from the first to the third egg collection time point in families M5 and M3 (Fig. 3 I, L). Family F1 produced only ten G_1_ progeny collected from two egg collections, but nine showed complete knock-in genotypes (Fig. 3 O). In family F4, knock-in events slightly decreased over time (Fig. 3 R). None of the analyzed individuals originating from the eGFP_gRNA2b injections carried the incomplete knock-in with only two bp changed instead of three that was observed with eGFP_gRNA2.

## Discussion

Genome editing in Medfly was successfully developed and evaluated via CRISPR-Cas HDR, using a short ssODN repair template to introduce point mutations in the eGFP marker gene of the transgenic line *TREhs43hid*^*Ala5*^_F1m2. We used two different gRNAs to target eGFP and one single-stranded repair template (ssODN_BFP) to achieve the conversion. After successful HDR, two mismatches were introduced to the target sequence of eGFP_gRNA2 (194C>G; 196T>C), while its PAM sequence remained intact. Regarding eGFP_gRNA2b, an HDR event introduced one mismatch to the target sequence (201C>G) and two to the PAM sequence (194C>G; 196T>C), whereby the PAM was eliminated (Fig. 1 A).

While only 50% of the injection survivors were fertile, we observed a very high efficiency of CRISPR-induced mutations, not only in the frequency of CRISPR-positive families (six out of seven fertile G_0_), but also in the penetrance within the families. Between 79 and 100% of G_1_ individuals within a family showed the phenotypical loss of eGFP fluorescence, indicating a mutation event and efficient targeting of the germ line in the G_0_ individuals. Sequence analysis confirmed these events and moreover revealed a knock-in rate of up to 96% (Fig. 3). We did not observe, however, the blue fluorescence that would be the phenotypic confirmation of a positive knock-in event. Reasons for this could be to the melanization of the medfly thorax or an autofluorescence overlapping with the spectrum of the ET DAPI BP filter.

Besides the three base pair BFP knock-in, we also detected a ‘partial knock-in’ with only two out of three base pairs changed when we used eGFP_gRNA2, but not with eGFP_gRNA2b. It was reported earlier that during HDR often only the part of the repair template actually overlapping with the deletion caused by the DSB is utilized [27, 33]. As small deletions are more common than large deletions, the probability for a mutation to be incorporated during the HDR event decreases with the increasing distance from the cleavage site. This finding could explain the missing third SNP in the first experiment (201C>G, ‘partial knock-in’), as that SNP is the one most distal to the DSB side of eGFP_gRNA2. However, we did not observe anything similar for eGFP_gRNA2b, although the distance between the cleavage site and the most distal SNP is similar (six bp for eGFP_gRNA2b, versus seven bp for eGFP_gRNA2). Alternatively, the occurrence of the partial knock-in could be the result of re-editing of the already modified locus [26], as the PAM of eGFP_gRNA2 remains intact after HDR whereas the PAM of eGFP_gRNA2b becomes eliminated. To ensure precise modification of the target site it is therefore important to include PAM-site mutations (silent) into the repair template [27].

This correlates also to the fact that the ‘complete knock-in’ increased over time in four out of six families. In contrast, the rate of the ‘partial knock-in’, which occurred in the two ‘eGFP_gRNA2’ families, decreased over time. This could indicate a general trend of increasing probability of knock-in events in egg collections from older adults. Such phenomenon paired with high efficiency would offer a possibility to save time and resources in mutagenic screens, especially while targeting genes which do not alter the phenotype. Further experiments will be needed, however, to corroborate these findings.

The additional use of Scr7 in the injection mix did not yield any phenotypic CRISPR events in Medfly. Scr7 inhibits DNA ligase IV, a key enzyme in the NHEJ pathway and has been shown to enhance the HDR rate in human cell cultures or mouse embryos [22]. Interestingly, the use of Scr7 increased the hatching rate compared to two injections without Scr7 (24.5% versus 6.6% and 5.1%, respectively). While injections of zinc finger nuclease together with circular donor DNA in ligase IV-deficient *Drosophila melanogaster* embryos successfully increased HDR compared to injections into WT flies [34, 35], Scr7, to our knowledge, has not been tested to enhance HDR in insects successfully. Therefore, further experiments with Scr7 at different concentrations will be interesting to investigate if it could have any effect in insects.

The use of an end-concentration of 300 mM KCl in the injection mix, as suggested previously for other Cas9 proteins [30, 36], seemed to help solubilizing the utilized Cas9-gRNA RNPs, as there were no issues regarding clogging of needles while injecting these concentrations of protein and RNA (360 ng/μl Cas9 protein and 200 ng/μl gRNA).

CRISPR-Cas allows a wide variety of genome editing strategies from small InDels at defined positions in the genome via NHEJ to the targeted introduction of point mutations (SNPs) via HDR all the way to the knock-out or knock-in of complete genes. While gene knock-in most probably will be classified as GMO, ‘non-traceable” CRISPR-induced mutations like InDels and SNPs, potentially might be regarded as non-GMO in the EU according to the ‘mutagenesis exemption’ foreseen in the GMO Directive [14, 15]. CRISPR-Cas can be another technique of mutagenesis and if part of the mutagenesis exemption of EU legislation in the future, it could be handled in a similar way to other mutagenesis techniques (e.g. UV or chemical mutagenesis) [15, 37]. In the US, the United States Department of Agriculture, Animal and Plant Health Inspection Service (USDA APHIS) recently classified CRISPR-edited organisms as ‘not regulated’ under *7 CFR part 340*. One example is the modified white button mushroom (*Agaricus bisporus*) with an anti-browning phenotype which is achieved by the introduction of small deletions (1-14 bp) in a specific polyphenol oxidase gene via CRISPR gene editing [38].

The classification of certain CRISPR-induced alterations as non-GMO would allow the application of this highly efficient and versatile technique for the development of new or improved strains for Medfly SIT programs and possibly for other related Tephritid fruit flies. The release of these strains would not be restricted via the GMO-Directive, and could be discussed in line with other solutions in terms of public acceptance, which is vital to the establishment and success of novel and safe pest control systems.

## Conclusions

Precise genome engineering is a powerful tool to improve and develop genetic pest control strategies to fight devastating and economically important pest species like the Mediterranean fruit fly. We demonstrate that genome editing via CRISPR-Cas HDR using a short, single-stranded DNA repair template is highly efficient in the Tephritid fruit pest *C. capitata*. If this high efficiency can be matched with larger repair templates remains to be seen. As there is the possibility that such ‘not-traceable’ CRISPR-induced mutations could be classified as non-GMO in the US as well as in Europe, the establishment of CRISPR-Cas genome editing in Medfly will be crucial for the development of new, genetically optimized strains for pest management systems like the classical SIT that are not restricted and GMO-free. This is the first report of successful CRISPR-Cas HDR genome editing in the family of Tephritidae, which contain a number economically important fruit pest species. The establishment of CRISPR-Cas genome editing in Medfly therefore is an important step towards the application of this technique to other Tephritid fruit pests like *Bactrocera dorsalis*, *B. oleae*, *Anastrepha ludens*, and *A. suspensa* and will be crucial for the development of non-transgenic and non-GM strategies to fight these pest insects.

## Material and methods

### Fly rearing

The *Ceratitis capitata* transgenic target line *TREhs43*^*Ala5*^_F1m2 flies [31] and wild type *Egypt-II* (*EgII*, obtained from the FAO/IAEA Agriculture and Biotechnology Laboratory, Seibersdorf, Austria) flies were maintained in a controlled environment (26°C, 48% RH, and a 14:10 light/dark cycle) and fed with a mixture of sugar and yeast extract (3v:1v), and water. Larvae were reared on a gel diet, containing carrot powder (120 g/l), agar (3 g/l), yeast extract (42 g/l), benzoic acid (4 g/l), HCl (25%, 5.75 ml /l) and Ethyl-4-hydroxybenzoat (2.86 g/l). Larvae and flies from injected embryos were reared under the same conditions. *TREhs43*^*Ala5*^_F1m2 flies used for CRISPR gene editing carry an eGFP marker under the control of the *D. melanogaster polyubiquitin* promotor [31]. The eGFP marker gene is expressed in head, thorax and legs of the adult fly. Flies were anesthetized with CO_2_ for screening, sexing, and the setup of backcrosses.

### CRISPR-Cas9 reagents

Purified Cas9 protein was obtained from PNA Bio Inc (catalog number CP01). The lyophilized pellet was reconstituted to a stock concentration of 1 μg/μl in 20 mM Hepes, 150 mM KCl, 2% sucrose and 1 mM DTT (pH 7.5) by adding 25 μl nuclease free H_2_O and stored at −80°C until use.

Linear double-stranded DNA templates for specific gRNAs were produced by a template-free PCR reaction with two partially overlapping oligos, containing 20 μl 5×Q5 reaction buffer, 10 μl dNTP Mix (2 mM each), 5 μl of each primer (10 μM) and 1 μl Q5 HF polymerase (2U) (New England Biolabs, NEB) in a total volume of 100 μl. PCR reactions were run in a Bio-Rad C1000 Touch thermal cycler [98°C, 30 s; 35 cycles of (98°C, 10 s; 58°C, 20 s; 72°C, 20 s); 72°C, 2 min] [39]. For the synthesis of the guide RNA eGFP_gRNA2 primers P_986 (GAAATTAATACGACTCACTATAGGCTCGTGACCACCCTGACCTAGTTTTAGAGCTAGAAATAGC) and P_369 (GCACCGACTCGGTGCCACTTTTTCAAGTTGATAACGGACTAGCCTTATTTTAACTTGCTATTTCTAGCTCTAAAAC) were used, for the synthesis of eGFP_gRNA2b primers P_1172 (GAAATTAATACGACTCACTATAGGCTGAAGCACTGCACGCCGTGTTTTAGAGCTAGAAATAGC) and P_369 were used. The forward primers (P_986, P_1172) encode the T7 polymerase-binding site followed by the specific gRNA target sequence and ending with the 20 nt complementary sequence that allows forward and reverse primers to anneal. Reverse primer (P_369) is a common oligonucleotide that can be used for all targets encoding the Cas9 interacting portion of the gRNA sequence [39]. The specific gRNA target sequence of gRNAs eGFP_gRNA2 and eGFP_gRNA 2b was previously described [28]. Size verification was carried out using 2 μl of the reaction while the remaining 98 μl were purified using a PCR purification kit (DNA Clean & Concentrator™-25; Zymo Research, Irvine, CA, USA) and eluted in 30μl elution buffer. Purity and concentration of the gRNA templates were measured with a spectrophotometer (BioTek Epoch2 microplate reader). gRNA *in vitro* transcription was performed with the HiScribe™ T7 High Yield RNA Synthesis Kit (NEB), using 500 ng purified DNA template for 16 h (overnight) at 37°C. RNA samples were treated with TURBODNase (Ambion, Oberursel, Germany) to remove possible DNA contamination, and purified using the MEGAclear purification kit (Ambion) [39]. Purified gRNAs were aliquoted and stored at −80°C until use.

The 140 bp single-stranded HDR template ‘ssODN_BFP’ (single-stranded oligodeoxynucleotide_blue fluorescent protein; P_1000_G/BFP_ssODN_Glaser) to convert eGFP into BFP was described previously [28] and synthetized by Eurofins Genomics (EXTREMer oligo, purified salt free, quality control by CGE). It differs from the eGFP sequence by 3 bases (194C>G, 196T>C, 201C>G), whereby the first change (194C>G; Thr65>Ser65) causes a reversion of eGFP back to wild-type GFP, the second (196T>C; Tyr66>His66) converts GFP to BFP. The third SNP (201C>G) is a silent mutation, to further reduce the target sequence similarity after HDR [28]. The sequence of ssODN_BFP was verified by sequencing (Macrogen Europe, Amsterdam), after performing PCR using Platinum Taq polymerase (Invitrogen), primers P_1160 (GGCATGGCGGACTTG) and P_1001 (CCTGAAGTTCATCTGCACCACC) in a Bio-Rad C1000 Touch thermal cycler [95°C, 2 min; 35 cycles of (95°C, 30 s; 50.5°C, 30 s; 72°C, 20 s); 72°C, 2 min]. PCR reaction contained 10 μl 10x Platinum PCR Buffer (-Mg), 1 μl MgSO_4_ 50 mM, 1 μl dNTP Mix (2 mM each), 1 μl of each primer (10 μM), 0.2 μl Platinum Taq polymerase and 440 ng DNA template in a total volume of 20 μl.

### Preparation of CRISPR injection mix

Injection mixes for microinjection of embryos contained 360 ng/μl Cas9 protein (1 μg/μl, PNA Bio, dissolved in its formulation buffer), 200 ng/μl gRNA_eGFP2 or gRNA_eGFP2b and 200 ng/μl ssODN_BFP in a 10 μl volume containing an end-concentration of 300 mM KCl, according to previous studies [30, 36, 40]. To inhibit NHEJ, 1 mM Scr7 (Xcess biosciences Inc., catalog number M60082-2) was added to the injection mix. All mixes were freshly prepared on ice followed by an incubation step for 10 min at 37°C to allow pre-assembly of gRNA-Cas9 ribonucleoprotein complexes and stored on ice prior to injections.

### Microinjection of embryos

For microinjection of homozygous *C. capitata TREhs43hid*^*Ala5*^_F1m2 embryos eggs were collected over a 45-90 min period. Eggs were prepared for injection as previously described [41] using chemical dechorionization (sodium hypochlorite, 3 min). In brief, embryos were fixed using double-sided sticky tape onto a microscope slide (Scotch 3M Double Sided Tape 665), and covered with halocarbon oil 700 (Sigma-Aldrich, Munich, Germany). Injections were performed using borosilicate needles (GB100F-10 with filaments; Science Products, Hofheim, Germany), drawn out on a Sutter P-2000 laser-based micropipette puller. The injection station consisted of a manual micromanipulator (MN-151, Narishige), an Eppendorf femtoJet 4i microinjector, and an Olympus SZX2-TTR microscope (SDF PLAPO 1×PF objective). The microscope slide with the injected embryos was placed in a Petri dish containing moist tissue paper in an oxygen chamber (max. 2 psi) and stored at 25°C, 60% RH for 72 hr to allow larval hatching. Hatched first instar larvae were transferred from the oil to larval food.

### Crossing and screening

Each G_0_ adult survivor was individually crossed to three *EgII* WT males or female virgins. Eggs were collected 3-4 times, with an interval of 1 to 2 days. Both G_0_ and G_1_ flies were screened for eGFP fluorescence phenotype to detect CRISPR mutagenesis events, G_1_ flies additionally were screened for BFP fluorescence.

### Genomic DNA extraction, PCR and sequencing

Genomic DNA was extracted from single G_1_ flies according to standard protocols. The DNA was used as template to amplify the region surrounding the gRNA target sites. PCR was performed in a 50 μl reaction volume using DreamTaq polymerase (Life Technologies), the eGFP-specific primers P_145 (ACTT AAT CGCCTTGCAGCACAT CC) and P_55 (TGTGATCGCGCTTCTCGTT), and 150 to 250 ng template in a Bio-Rad C1000 Touch thermal cycler [95°C, 3 min; 35 cycles of (95°C, 30 s; 58°C, 30 s; 72°C, 1 min); 72°C, 5 min]. Afterwards, the size of the PCR product was verified by running an aliquot of the reaction on an agarose gel. The remaining PCR product was purified using a PCR purification kit (DNA Clean & Concentrator™-25; Zymo Research). All PCR products were verified by sequencing (Macrogen Europe, Amsterdam; with Primer P_145).

### Verification of mutations and off-target assessment

Assessment of potential off-targets effects of gRNA2 and gRNA2b was performed using the *C. ceratitis* genome annotation Ccap 1.1 (GCF_000347755.2_Ccap_1.1_genomic.fna.gz) from NCBI also using Geneious [42]. Verification of CRISPR-induced mutations on sequencing results was performed using the Software Package Geneious 10.2.2 [43] by mapping the sequencing results of G_1_ individuals to the eGFP reference sequence [31].

### Equipment and settings for screening and image acquisition

Screening of transgenic flies was performed using a Leica M165 FC stereo microscope with the PLAN 0.8× LWD objective and the following epifluorescence filters: GFP-LP (Excitation 425/60 nm barrier 480 LP nm), YFP (excitation 510/20 nm; barrier 560/40 nm) or ET DAPI BP (excitation 395/25 nm; barrier 460/50 nm). For bright field and fluorescent image acquisition of living flies (anesthetized with CO_2_ and placed on a 4°C cooler) was carried out using a fully automated Leica M205FC stereo microscope with a PLANAPO 1.0× objective and a 1× Leica DFC7000 T camera using Leica LAS X software. In order to enhance screen and print display of the pictures the image processing software Adobe Photoshop CS5.1 was used to apply moderate changes to image brightness and contrast. Changes were applied equally across the entire image and throughout all images.

## Availability of data and material

All data generated or analyzed are included in this article.

## Authors’ contributions

RA performed research; MFS, RA and IH designed research; RA, MFS and IH analyzed data; and RA, IH and MFS wrote the paper. MFS and IH were group leaders for the project. All authors have read and approved the final version of the manuscript.

## Acknowledgements

We wish to thank Tanja Rehling, Derya Arslan and Julia Hehner for technical assistance, and Kate Sutton for proofreading of the manuscript.

